# Predicting factors for long-term survival in patients with out-of-hospital cardiac arrest – a propensity score-matched analysis

**DOI:** 10.1101/664441

**Authors:** Anna Lena Lahmann, Dario Bongiovanni, Anna Berkefeld, Maximilian Kettern, Lucas Martinez, Rainer Okrojek, Petra Hoppmann, Karl-Ludwig Laugwitz, Markus Kasel, Salvatore Cassese, Robert Byrne, Sebastian Kufner, Erion Xhepa, Heribert Schunkert, Adnan Kastrati, Michael Joner

**Affiliations:** Department of cardiology, German Heart Center Munich, Technical University Munich, Munich, Germany; Department of cardiology, Klinikum rechts der Isar, Technical University Munich, Munichh, Germany; DZHK (German Center for Cardiovascular Research), partner site Munich Heart Alliance, Munich, Germany

## Abstract

**Background:** Out-of-hospital cardiac arrest (OHCA) is one of the leading causes of death worldwide, with acute coronary syndromes accounting for most of the cases.

While the benefit of early revascularization has been clearly demonstrated in patients with ST-segment-elevation myocardial infarction (STEMI), diagnostic pathways remain unclear in the absence of STEMI. We aimed to characterize OHCA patients presenting to 2 tertiary cardiology centers and identify predicting factors associated with survival.

**Methods:** We retrospectively analyzed 519 patients after OHCA from February 2003 to December 2017 at 2 centers in Munich, Germany. Patients undergoing immediate coronary angiography (CAG) were compared to those without. Propensity score (PS) matching analysis and multivariate regression analysis were performed to identify predictors for improved outcome.

**Results:** Immediate CAG was performed in 385 (74.1%) patients after OHCA with presumed cardiac cause of arrest.

As a result of multivariate analysis after propensity score matching, we found that ROSC at admission and immediate CAG were associated with better 30-days-survival [(OR, 6.54; 95% CI, 2.03-21.02), (OR, 2.41; 95% CI, 1.04-5.55)], and 1-year-survival [(OR, 4.49; 95% CI, 1.55-12.98), (OR, 2.54; 95% CI, 1.06-6.09)].

**Conclusions:** In our study, ROSC at admission and immediate CAG were independent predictors of survival in cardiac arrest survivors. Improvement in prehospital management including bystander CPR and best practice post-resuscitation care with optimized triage of patients to an early invasive strategy may help ameliorate overall outcome of this critically-ill patient population.

## Introduction

Out-of-hospital cardiac arrest (OHCA) is a leading cause of death in western countries and still associated with poor prognosis despite improvement in emergency care and post-resuscitation management in recent years [1–4].

Increased rates of bystander resuscitation (CPR), and widespread availability of automated external defibrillators (AEDs) substantially contributed to improved patient outcome; in addition, it has been shown that targeted temperature management and early revascularization improve survival of patients with cardiac arrest caused by myocardial infarction [5–7].

Contemporary management of patients with ST-segment elevation recommends immediate coronary angiography (CAG) after cardiac arrest, including ad-hoc percutaneous intervention (PCI) if necessary. Current guidelines also suggest to consider immediate CAG for those patients with presumed cardiac cause of arrest in the absence of STEMI owing to the high incidence of coronary artery disease in these patients [8–9].

In this context, it remains challenging to select candidates for early CAG given the extremely heterogeneous population of cardiac arrest survivors and the difficulty to retrieve etiologic information in this specific setting [8,10].

The aim of this study was to characterize patients presenting to 2 major tertiary cardiology centers with intensive care capability after OHCA and stratification into those undergoing immediate coronary angiography or not. Furthermore, it was our goal to identify independent prognostic factors associated with survival in patients with OHCA.

## Materials and Methods

We collected retrospective data of patients after out-of-hospital cardiac arrest from February 2003 to December 2017 at two centers in Munich, Germany. OHCA data were collected according to the Utstein recommendations and the ethics committee of the Technical University Munich (approval number 343/17 S) waived informed consent due to the observational nature of the study.^11^

### Endpoint definitions

The primary outcome was survival at 30 days and one year, which was assessed by medical records or by telephone interview of the attending physicians; secondary outcome was functional status at discharge which was evaluated using the Cerebral Performance Category (CPC) score.^12^

### Patient flow

Triage of patients to undergo immediate CAG was left to the responsible physician’s discretion. When available, paramedical information, ECG findings and echocardiographic abnormalities aided in decision-making. An immediate coronary angiogram and subsequent PCI was performed if necessary, using standard techniques. Patients were subsequently admitted to the intensive care unit (ICU) for standard post-resuscitation care including targeted temperature management if indicated.

Immediate CAG was defined as coronary angiography performed within two hours after admission to hospital.

A coronary lesion resulting in >75% reduction of luminal diameter by visual estimation was considered significant and PCI was deemed successful when resulting in a residual stenosis of <30%.

Culprit lesion morphology was determined by angiography and defined as acute coronary occlusion, presence of thrombus, severe narrowing in the presence or absence of thrombus, and unstable-appearing lesions with high likelihood to trigger ischemia responsible for cardiac arrest.

STEMI and NSTEMI, as well as the other causes of cardiac arrest, were determined by review of ECGs after return to spontaneous circulation (ROSC) and by review of patients’ charts including CAG and serum parameters.

### Statistical analysis

Continuous variables are presented as mean ± SD. Distribution of data was checked for normality using the Shapiro-Wilk goodness-of-fit test and differences analyzed with the Wilcoxon-signed rank sum test in case of non-parametric data and student’s t-test in the event of normal distribution. Categorical variables are presented as number (%) and were analyzed using the χ^2^-test. Univariate logistic regression analysis was performed to derive crude 30-days and 1-year-survival rates among patients undergoing immediate CAG or not.

To minimize selection bias of patients undergoing immediate CAG and to control for potential confounding factors, we conducted propensity score (PS) regression analysis and generated two matching cohorts of patients undergoing immediate CAG or not. Matching variables were hypercholesterolemia, daytime presentation, witnessed arrest, arrest at home, active smoker, former smoker, ROSC at admission and female. All available univariate factors were subsequently entered into a multivariate generalized linear regression model. Selection of covariates in the multivariate regression model was performed using the LASSO (Least Absolute Shrinkage and Selection Operator) regression method after entering all available candidates. These variables were asystole in the initial ECG, ROSC at admission, bystander CPR, daytime presentation, witnessed arrest, active smoker, former smoker, diabetes, hypercholesterolemia, family history, female, history of prior myocardial infarction, history of coronary artery disease, arterial hypertension and immediate CAG.

Parameters achieving p-values <0.05 were considered statistically significant and odds ratios derived with 95% confidence intervals. Analysis was performed using JMP Pro (software version 13.0, Cary, NC, USA) and SPSS (software version 22 with FUZZY extension bundle, IBM, Armonk, NY, USA).

## Results

We analyzed a total of 519 patients with out-of-hospital cardiac arrest between February 2003 and December 2017 that were consecutively admitted to one of two tertiary centers in Munich, Germany. The baseline demographic characteristics are shown in (Table 1).

**Table 1.** Baseline characteristics of patients with and without immediate CAG before and after propensity score analysis.

| Entire population |  |  |  | Matched population |  |  |
| --- | --- | --- | --- | --- | --- | --- |
|  | Immediate<br>CAG<br>n=385 | No immediate<br>CAG<br>n=134 | p-value | Immediate<br>CAG<br>n=87 | No immediate CAG<br>n=88 | p-value |
| Female | 89/385 (23.1) | 45/134 (33.6) | **0.02** | 21/87 (24.1) | 23/88 (26.1) | 0.76 |
| Age | 65.1±13 | 66.9±16 | 0.11 | 65 ± 13 | 67 ± 16 | 0.76 |
| Former smoker | 143/326 (43.9) | 25/83 (30.1) | **0.02** | 19/64 (29.7) | 21/68 (30.9) | 0.88 |
| Diabetes | 79/342 (23.1) | 25/87 (28.7) | 0.23 | 19/71 (26.8) | 20/67 (29.9) | 0.69 |
| Hypercholesterolemi<br>a | 208/327 (63.6) | 32/82 (39.0) | **<0.001** | 27/61 (44.3) | 26/65 (40.0) | 0.63 |
| Hypertension | 247/330 (74.9) | 64/90 (71.1) | 0.47 | 47/64 (73.4) | 51/71 (71.8) | 0.83 |
| Family history | 48/321 (15.0) | 7/78 (9.0) | 0.17 | 11/62 (17.7) | 6/64 (9.4) | 0.17 |
| History of coronary<br>artery disease | 111/369 (30.0) | 29/98 (29.6) | 0.93 | 19/80 (23.8) | 25/76 (32.9) | 0.20 |
| History of myocardial<br>infarction | 68/361 (18.8) | 18/96 (18.8) | 0.98 | 11/75 (14.7) | 15/73 (20.6) | 0.35 |

Data are presented as n (%) of the total cohort

ECG at first contact with emergency medical service (EMS) showed ventricular fibrillation in 233/376 (62.0%) of the patients who underwent immediate CAG, and in 26/125 (20.8%) in the no immediate CAG group (Table 2) as most frequent initial rhythm.

**Table 2.** Patients with and without immediate CAG according to initial ECG before and after propensity score analysis.

| Entire population |  |  |  | Matched population |  |  |
| --- | --- | --- | --- | --- | --- | --- |
|  | Immediate<br>CAG<br>n=385 | No immediate<br>CAG<br>n=134 | p-value | Immediate CAG<br>n=87 | No immediate<br>CAG<br>n=88 | p-value |
| Asystole | 89/376 (23.7) | 66/125 (52.8) | <0.001 | 24/85 (28.2) | 38/81 (46.9) | 0.01 |
| AV-Block | 3/376 (0.8) | 0/125 (0.0) | 0.32 | 0/85 (0.0) | 0/81 (0.0) | 0.0 |
| Bradycardia | 9/376 (2.4) | 1/125 (0.8) | 0.27 | 1/85 (1.2) | 0/81 (0.0) | 0.33 |
| VT | 8/376 (2.1) | 3/125 (2.4) | 0.86 | 0/85 (0.0) | 2/81 (2.5) | 0.15 |
| VFib | 233/376 (62.0) | 26/125 (20.8) | <0.001 | 51/85 (60.0) | 21/81 (25.9) | <0.001 |
| PEA | 22/376 (5.9) | 22/125 (17.6) | <0.001 | 5/85 (5.9) | 15/81 (18.5) | 0.01 |
| Miscellaneous | 12/376 (3.2) | 7/125 (5.6) | 0.22 | 4/85 (4.7) | 5/81 (6.2) | 0.68 |

Data are presented as % of the total cohort. Abbreviations: VT, ventricular tachycardia; VFib, ventricular fibrillation; PEA, pulseless electrical activity.

Major causes of cardiac arrest were STEMI and NSTEMI in 111/517 (21.5%) and 141/517 (27.3%) of all patients, respectively (Table 3).

**Table 3.**
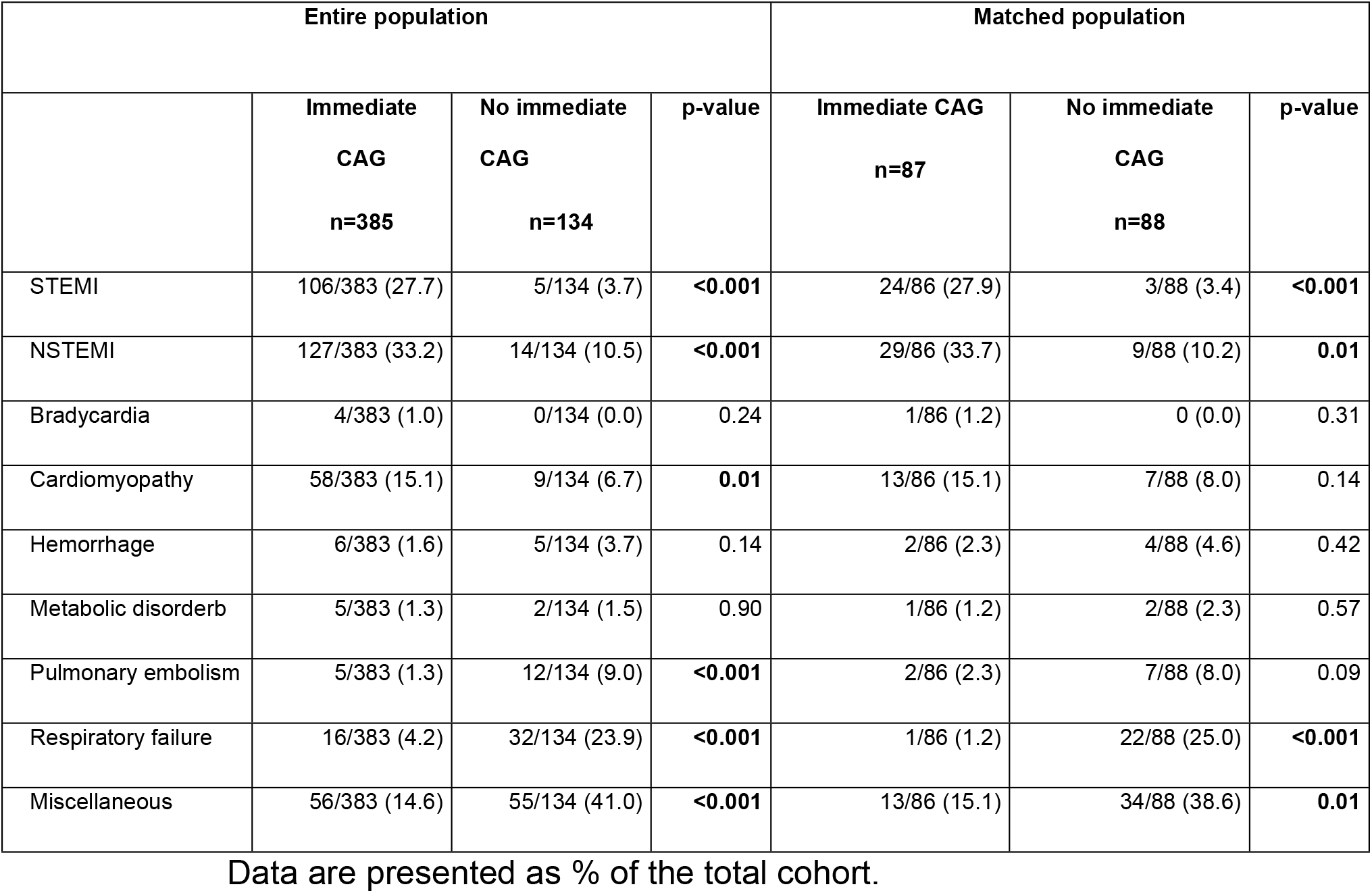
Cause of cardiac arrest in patients with and without immediate CAG before and after propensity score analysis.

In the 385 patients undergoing immediate CAG, a culprit lesion was identified in 247/385 (64.2%) (Fig 1A and 1B).

**Figure 1A+1B.**
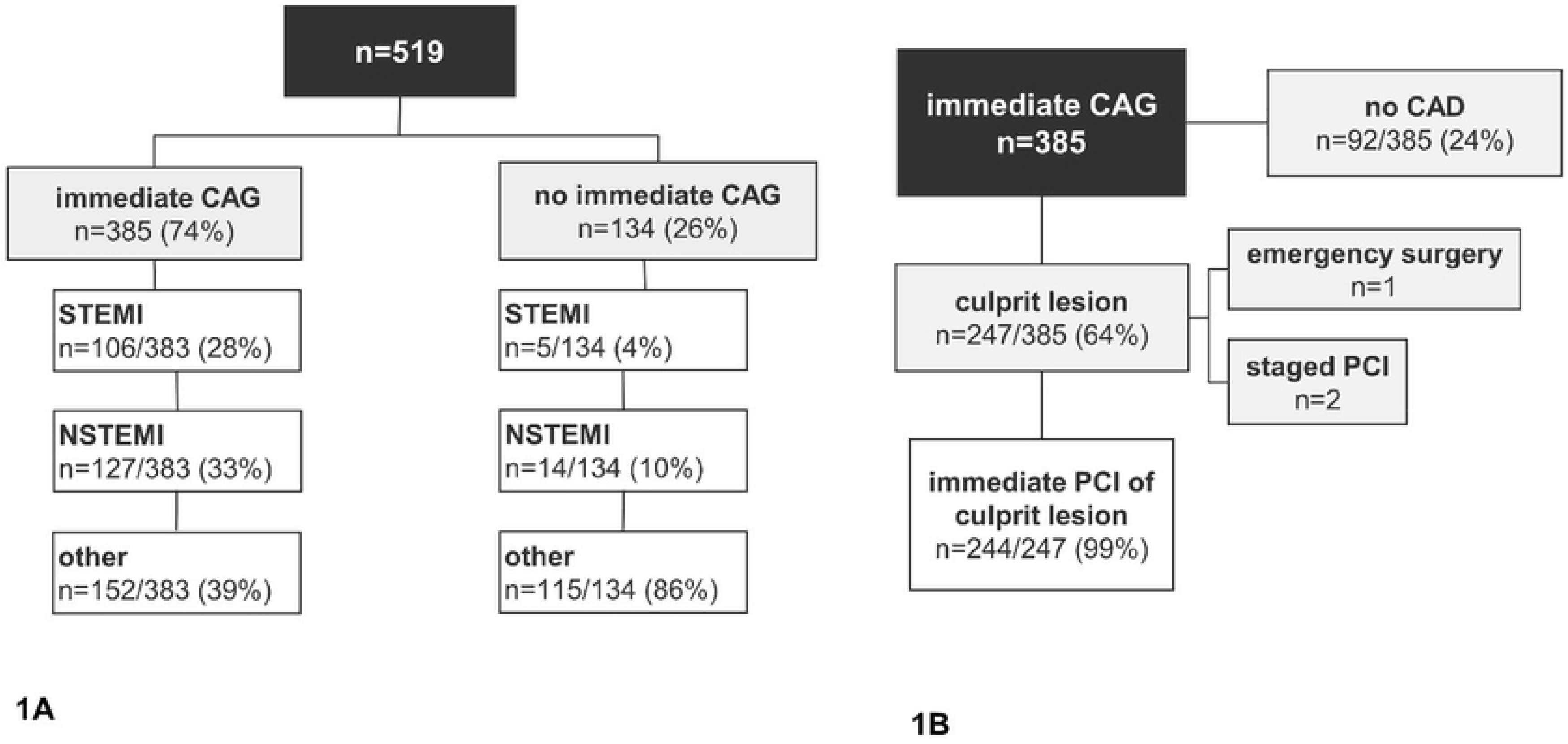
Patient flow diagrams. With regards to pre-hospital management, 336/479 (70.1) of the patients received bystander CPR and the arrest was witnessed in 381/487 (78.2). Arrest at home was found in 298/507 (58.8) of the patients. Mean time to ROSC was 14.4 ± 11.3 minutes in the immediate CAG group and 12.9 ± 11.6 in patients without with no significant difference between patients undergoing immediate CAG or not (Table 4).

**Table 4.** Preclinical and hospital data of patients with and without immediate CAG after propensity score analysis.

| Entire population |  |  |  |  | Matched population |  |
| --- | --- | --- | --- | --- | --- | --- |
|  | Immediate<br>CAG<br>n=385 | No<br>immediate<br>CAG<br>n=134 | p-value | Immediate CAG<br>n=87 | No<br>immediate<br>CAG<br>n=88 | p-value |
| **Arrest data** |  |  |  |  |  |  |
| Bystander CPR | 259/355<br>(73.0) | 77/124 (62.1) | **0.02** | 55/80 (68.8) | 53/82 (64.6) | 0.58 |
| ROSC at admission | 327/384<br>(85.2) | 86/133 (64.7) | **<0.001** | 66/87 (75.9) | 63/88 (71.6) | 0.52 |
| Time to ROSC<br>(min) | 14.4 ± 11.3 | 12.9 ± 11.6 | 0.23 | 18.7 ± 14.9 | 14.1 ± 12.4 | 0.07 |
| Daytime<br>presentation | 330/384<br>(85.9) | 106/131 (80.9) | 0.17 | 73/87 (83.9) | 78/88 (88.6) | 0.36 |
| Witnessed arrest | 295/364<br>(81.0) | 86/123 (69.9) | **0.01** | 68/82 (82.9) | 68/85 (80.0) | 0.63 |
| Arrest at home | 207/380<br>(54.5) | 91/127 (71.7) | **0.01** | 61/85 (71.8) | 60/87 (69.0) | 0.69 |
| **Hospital data** |  |  |  |  |  |  |
| pH-value in first<br>BGA | 7.1 ± 0.2 | 7.0 ± 0.2 | **<0.001** | 7.09 ± 0.24 | 7.02 ± 0.21 | **0.03** |
| pO<sub>2</sub> in first BGA<br>(mmHg) | 191.8 ± 145.2 | 168.0 ± 137.3 | 0.08 | 224.4 ± 172.9 | 170.6 ± 142.8 | 0.14 |
| Lactate in first BGA<br>(mmol/l) | 8.8 ± 4.8 | 12.4 ± 8.2 | **<0.001** | 9.6 ± 5.2 | 11.5 ± 6.6 | 0.17 |
| NSE after 48 hours<br>(ng/ml) | 74.4 ± 104.3 | 127.9 ± 145.0 | 0.06 | 92.5 ± 127.6 | 98.4 ± 112.8 | 0.68 |

Data are presented as n (%) or as mean ± SD unless indicated otherwise. Abbreviations: CAG, coronary angiography; ROSC, return of
spontaneous circulation; CPR, cardiopulmonary resuscitation; EMS, emergency medical service; BGA, blood gas analysis; NSE, neuron-specific enolase; CPC, cerebral performance category; TTM, targeted temperature management.

Coronary angiographic and intervention findings are shown in Table 5.

**Table 5.** Coronary angiographic and intervention findings in patients with immediate CAG before and after propensity score analysis.

| Entire population<br>n=385 |  | Matched population<br>n=88 |
| --- | --- | --- |
| Single-vessel disease | 64/385 (16.6) | 21/87 (24.1) |
| Double-vessel disease | 81/385 (21.0) | 17/87 (20.0) |
| Triple-vessel disease | 148/385 (38.4) | 27/87 (31.0) |
| Overall culprit lesions | 247/385 (64.2) | 58/87 (66.7) |
| Culprit lesion LAD | 115/247 (46.6) | 32/87 (36.8) |
| Culprit lesion LCx | 45/247 (18.2) | 9/87 (10.3) |
| Culprit lesion RCA | 87/247 (35.2) | 17/87 (19.5) |
| Immediate PCI of culprit lesion | 244/385 (63.4) | 57/87 (65.5) |
| PCI of more than culprit lesion | 32/385 (8.3) | 6/87 (6.9) |
| Other significant lesions<br>not treated in the emergency setting | 62/384 (16.2) | 11/87 (12.6) |

Data are presented as n (%) or as mean ± SD. Abbreviations: EF, ejection fraction; LAD, left anterior descending artery; LCx, left
circumflex artery; RCA, right coronary artery; PCI, percutaneous coronary intervention; CAD, coronary artery disease.

With regards to pre-hospital management, 336/479 (70.1) of the patients received bystander CPR and the arrest was witnessed in 381/487 (78.2). Arrest at home was found in 298/507 (58.8) of the patients. Mean time to ROSC was 14.4 ± 11.3 minutes in the immediate CAG group and 12.9 ± 11.6 in patients without with no significant difference between patients undergoing immediate CAG or not (Table 4).

Patients in the immediate CAG group had significant higher pH-values and lower lactate levels in the first blood gas analysis (7.11 ± 0.20 vs. 7.00 ± 0.21, p<0.001 and 8.8 ± 4.8 mmol/l vs. 12.4 ± 8.2 mmol/l, p<0.001).

Overall 30-days-survival of the 519 patients that were included in the study was 50.1% compared to a 1-year-survival of 37.6%. 30-days-survival as well as survival after one year was significantly better in the immediate CAG group [221/370 (59.7) vs. 30/131 (22.9), p<0.001] and [161/341 (47.2) vs. 14/124 (11.3), p> 0.001] than in patients without.

Neurological function was evaluated at hospital discharge. In our cohort, good neurological function (CPC 1&2) was found in 164/299 patients (54.8%) at discharge (Table 6).

**Table 6.** Cerebral Performance Category Score 1&2 at discharge in patients with immediate and no immediate CAG before and after propensity score analysis.

| Entire population |  |  | Matched population |  |  |
| --- | --- | --- | --- | --- | --- |
| Immediate CAG<br>n=385 | No immediate<br>CAG<br>n=134 | p-value | Immediate CAG<br>n=87 | No immediate CAG<br>n=88 | p-value |
| 151/261 (57.9) | 13/38 (34.2) | **0.01** | 23/50 (46.0) | 10/28 (35.7) | 0.38 |

Data are presented as n (%). Abbreviations: CPC, cerebral performance category.

### Matched population

After applying PS matching to reduce confounding factors arising from differential selection of patients undergoing immediate CAG or not, two comparable cohorts were generated to identify predicting factors associated with survival after OHCA.

By multivariate analysis, we found ROSC at admission (OR, 6.54; 95%CI, 2.03-21.02) and immediate CAG (OR, 2.41; 95%CI, 1.04-5.55) were predictors for better 30-days-survival (Fig 2) and 1-year-survival [(OR, 4.49; 95%CI, 1.55-12.98) (OR, 2.54; 95%CI, 1.06-6.09)] (Fig 3).

**Figure 2.**
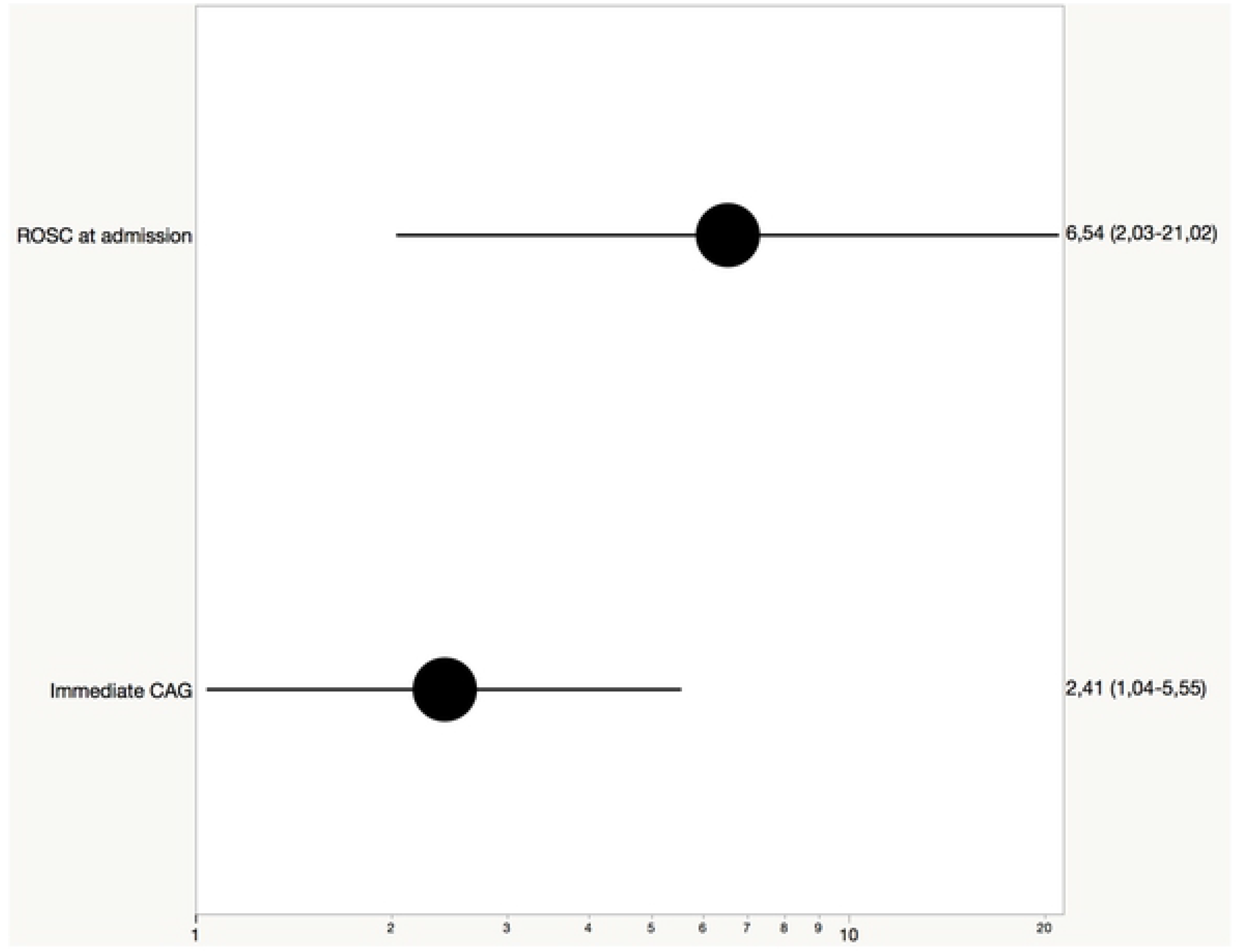
Forest plot with Odds ratios and 95% confidence intervals of factors associated with 30-days-survival after propensity score-matched analysis.

**Figure 3.**
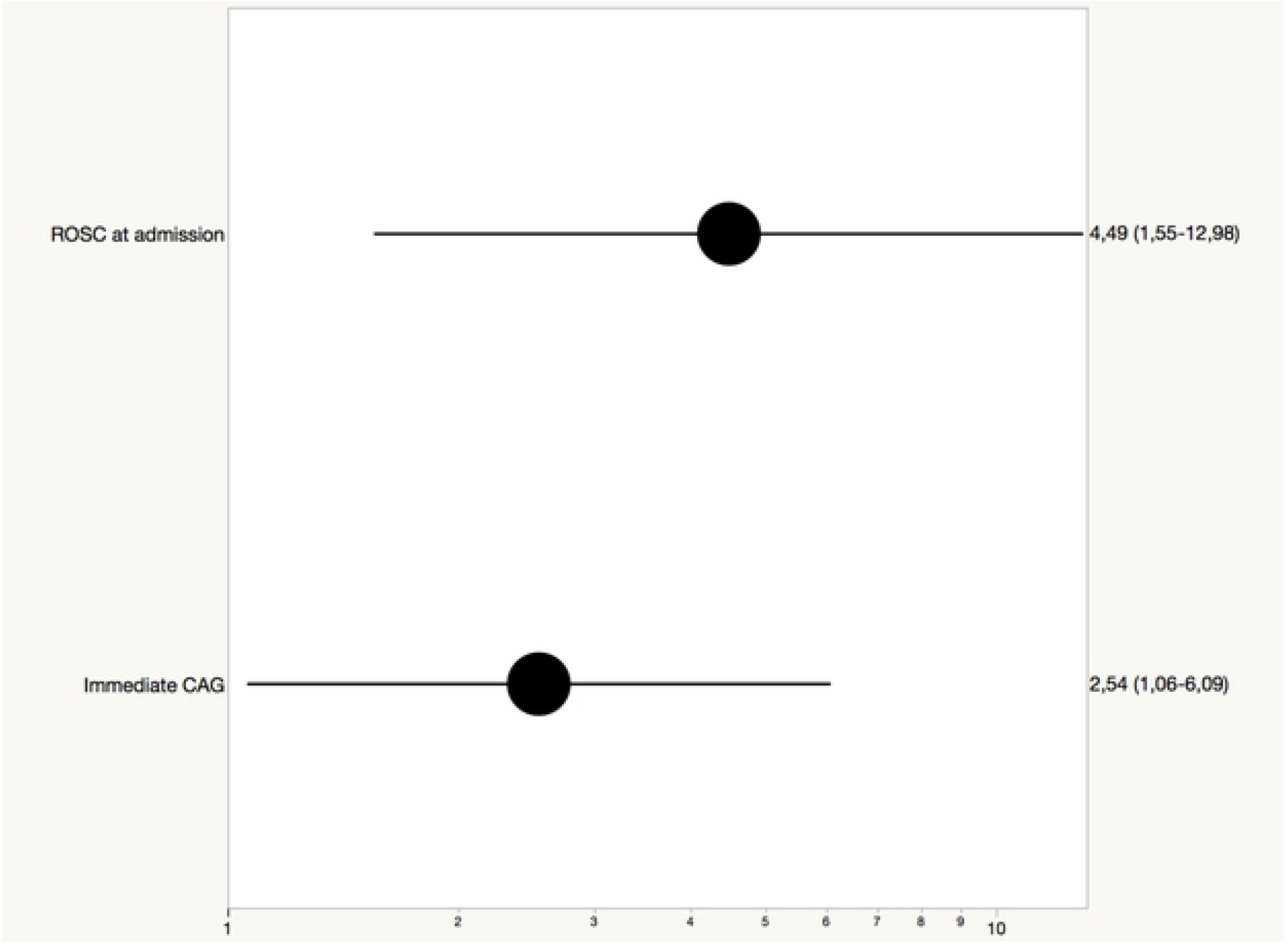
Forest plot with Odds ratios and 95% confidence intervals of factors associated with 1-year-survival after propensity score-matched analysis.

## Discussion

Out-of-hospital cardiac arrest (OHCA) is a leading cause of death in developed countries. Both, resuscitation and intensive care management of patients after OHCA have notably improved over the recent years [6,8,13,14].

Although OHCA is mostly caused by acute myocardial infarction, it is unknown whether early coronary angiography is associated with improved survival in all patients after OHCA.

Our study represents one of the largest cohorts analyzing patients after out-of-hospital arrest with long-term survival and coronary angiographic data, collected at 2 centers specialized in coronary intervention and post-resuscitation care. It was our aim to characterize this extremely heterogeneous and critically ill population especially with regards to the impact of immediate coronary angiography. In this regard, the most salient findings can be described as follows:

→ Of the 519 patients admitted to the 2 participating centers, 74.1% of the study population underwent emergency coronary angiography.
→ 233/383 patients (60.8%) in the immediate CAG group were diagnosed with STEMI (n=106) or NSTEMI (n=127) after having completed diagnostic work-up including coronary angiography versus 19/134 patients (14.2%) in the no immediate CAG group.
→ In our cohort, good neurological function at discharge, defined as CPC 1 or 2, was achieved in 164/299 (54.8%)
→ 30-days-survival and survival after one year was significantly higher among patients undergoing immediate CAG compared to patients without immediate CAG [221/370 (59.7) vs. 30/131 (22.9), p<0.001] and [161/341 (47.2) vs. 14/124 (11.3), p> 0.001]
→ In our cohort, good neurological function at discharge, defined as CPC 1 or 2, was achieved in 164/299 (54.8%)
→ By multivariate analysis after PS matching, we identified ROSC at admission and immediate CAG as independent predictive factors for 30-days-survival and 1-year-survival in patients with OHCA

### Influence of study population and immediate coronary angiography on survival

Our study is composed of patients presenting to 2 tertiary centers specialized in cardiac care with capability for primary PCI and post-resuscitation care after OHCA. Consequently, pre-selection of patients by on-site emergency physicians and paramedical staff is very likely to impact significantly on overall findings and survival [15]. Additional selection bias arises from early triage after patient admission, where those with ST-segment elevation and other ECG abnormalities suggestive of ischemia are likely to undergo immediate CAG. To reduce these confounding factors, we performed propensity score regression analysis, which confirmed the survival benefit of patients undergoing immediate CAG. Along these lines, the overall survival rate of 50.1% in our study further supports the previously encountered phenomenon that triage into early angiographic evaluation in dedicated cardiac care centers may help improve outcome of these critically ill patients.

Previous studies suggested a high incidence of coronary artery disease in patients without obvious extracardiac cause of arrest, proposing early coronary angiography to be performed in most patients [8,16,17].

Corroborated by our findings, these results emphasize the relevance of appropriate patient selection for an invasive diagnostic strategy [8,18].

### Factors impacting on neurological outcome and survival

Good neurological function (defined as CPC 1&2) at discharge was achieved in 164/299 (54.8) of the patients, comparable to previous studies [19–21], with improved neurological outcome for patients undergoing immediate CAG. While previous studies addressing this association have already suggested a favorable effect of early angiographic assessment in post-resuscitation care, most, if not all studies including ours are hampered by non-randomized and retrospective design, increasing the chance to receive immediate CAG in patients with obvious ECG abnormalities and favorable OHCA resuscitation response including those with early ROSC, bystander CPR and witnessed arrest. Subsequently, patients undergoing immediate CAG are at higher likelihood for a favorable neurological outcome compared to those patients where early triage during patient presentation is influenced by unfavorable OHCA resuscitation response. Nevertheless, we and other authors have shown that culprit coronary lesions are detected in a large proportion of patients undergoing immediate CAG providing an opportunity to improve neurological outcome by primary PCI and by using propensity matching, we aimed to reduce the inherent selection bias of observational studies.

Patients with immediate CAG had significantly more often bystander CPR and a witnessed arrest, which is consistent with more shockable rhythms in this group leading to ROSC at admission, which we have shown to be predictive of favorable outcome [22,23]. In line with this, our findings are congruent with those of Stiell et al. in such that bystander resuscitation is a major factor of survival and neurological outcome after cardiac arrest. In addition, the wide-spread availability of automated external defibrillators (AEDs) has recently been shown to be associated with favorable outcome, which highlights the importance of prehospital resuscitation quality [14,24].

In our study, we observed an overall survival rate of 50.1%, which is in agreement with a priori selection of best candidates for early CAG. Furthermore, it was performed at 2 centers, where prehospital management of OHCA is performed according to standardized protocols with great experience in the treatment of acute coronary syndrome and cardiac arrest, which might have contributed to better survival rates than elsewhere and was recently pointed out by Soholm et al [15].

In our cohort, 30-days-survival was greater among patients with immediate coronary angiography compared to those without. These results are similar to those shown in the meta-analysis by Camuglia et al. with the limitation that they only considered survival to hospital discharge [21]. Furthermore, whether immediate CAG with subsequent PCI is associated with improved outcome or whether comorbidity and yet unidentified factors prevail to determine outcome in this critically-ill patient population remains to be investigated in dedicated prospective trials.

### Limitations

Our observations are obviously limited by the non-randomized, retrospective design of the study. Outcomes in the heterogenous population of patients after OHCA are likely impacted by selection and best practice of treating physicians.

## Conclusions

Our findings support that triage for immediate coronary angiography as part of post-resuscitation care facilitated by rapid interdisciplinary decision-making is of major importance. Furthermore, we confirmed the favorable impact of optimal prehospital management with improved outcome after witnessed arrest probably resulting in ROSC at admission.

Immediate coronary angiography in cardiac arrest survivors appears to be associated with improved survival and may enable therapeutic algorithms, particularly identifying those who may benefit from acute revascularization therapy.

